# Breaking the spell of nestedness

**DOI:** 10.1101/216564

**Authors:** Clàudia Payrató Borrás, Laura Hernández, Yamir Moreno

**Affiliations:** Laboratoire de Physique Théorique et Modélisation, UMR CNRS, Université de Cergy-Pontoise, 2 Avenue Adolphe Chauvin, F-95302, Cergy-Pontoise Cedex, France.; Institute for Biocomputation and Physics of Complex Systems (BIFI), University of Zaragoza, Spain.; Department of Theoretical Physics, Faculty of Sciences, University of Zaragoza, Spain.; Complex Networks and Systems Lagrange Lab, Institute for Scientific Interchange, Turin, Italy.; Complexity Science Hub Vienna, Austria.

**Keywords:** Mutualistic Ecosystems, Nestedness, Exponential Random Graph, Null Models

## Abstract

Mutualistic interactions, which are beneficial for both interacting species, are recurrently present in ecosystems. Observations of natural systems showed that, if we draw mutualistic relationships as binary links between species, the resulting bipartite network of interactions displays a widespread particular ordering called nestedness [1]. On the other hand, theoretical works have shown that a nested structure has a positive impact on a number of relevant features ranging from species coexistence [2], to a higher structural stability of communities and biodiversity [3,4]. However, how nestedness emerges and what are its determinants, are still open challenges that have led to multiple debates to date [5–7]. Here, we show, by applying a theoretical approach to the analysis of 167 real mutualistic networks, that nestedness is not an irreducible feature, but a consequence of the degree sequences of both guilds of the mutualistic network. Remarkably, we find that an outstanding majority of the analyzed networks does not show statistical significant nestedness. These findings point to the need of revising previous claims about the role of nestedness and might contribute to expand our understanding of how evolution shapes mutualistic interactions and communities by placing the focus on the local properties rather than on global quantities.

## Introduction

Nested patterns are ubiquitous in ecological systems. This observation has triggered an intense research aimed at defining and measuring nestedness [8] as well as at explaining its origin [9–11]. The interest in deciding whether a system is nested or not goes beyond characterizing it from a merely topological viewpoint. Admittedly, it has been suggested that nestedness plays an important role in biodiversity persistence, a claim which is nevertheless the subject of an ongoing and intense debate [2–4]. Furthermore, the relevance of nestedness as a suitable indicator to characterize mutualistic ecosystems has been recently challenged [12,13]. Several works have proposed alternative properties of the observed networks to link relevant structural characteristics with the system’s dynamics, in particular, the networks’ assortativity or the heterogeneity in the number of interactions of the species [13–15].

A key question is then whether nestedness, conceived as a global trait of the emerging architecture, is actually relevant and informative of the ecosystems’ dynamics, or contrarily, it just derives from lower order properties of the interaction network. In addition, elucidating the previous question would also allow to solve another open challenge, namely, which is the right *null model* against which one should assess nestedness? The latter is a relevant issue by itself, as any claim concerning statistical significance of a nested pattern implicitly involves its comparison with a null hypothesis (model). In order to address the aforementioned questions, we analyze an empirical set of 167 mutualistic networks (see Methods) to determine if, indeed, the observed amount of nestedness in real ecosystems could solely arise from the empirical degree sequences. Our choice is rooted on a a theoretical work [16] that showed that the geometric curve that delimits the region with interactions in an ideally nested matrix [17] can be ultimately related, by means of an approximation, to the *degree distributions* of both guilds of the corresponding bipartite network.

We constructed a grand canonical ensemble for each empirical ecological web under the constraint that, for the two guilds, the degree sequences in the ensemble match on *average* the empirical ones (see *Methods* and the *SM*). This methodological approach has the advantage that possible missing links or overrated interactions, that have been suggested to lead to impoverished ecological data [18], are dealt with in a proper way. In fact, constraining the randomized degree sequences to be equivalent to the empirical ones only on average limits the possible effects of noisy data, while assuring that results are not dependent on specific details. At variance with previous works that have imposed similar constraining rules [19], here we apply a recently introduced randomizing scheme [20,21] that treats the ensemble from a statistical physics perspective, yielding the maximum entropy network ensemble such that the degree sequence of the empirical network is found with maximum likehood (see *Methods*). This enforces, as aimed, the ensemble’s mean degree sequences to be the empirical ones whilst precluding common biases of other sampling techniques [22], and allows us to obtain the probability that two potential partners interact in the randomized ensemble, see Fig.1. Moreover, one can explicitly write expressions for the main statistical moments of any network property that can be analytically formulated in terms of the elements of the bipartite adjacency matrix. Since the well-known NODF metric for nestedness fulfills such a condition, we have derived the analytical expressions for the mean and the standard deviation of the nestedness’ distribution, see *Methods*.

**Figure 1.**
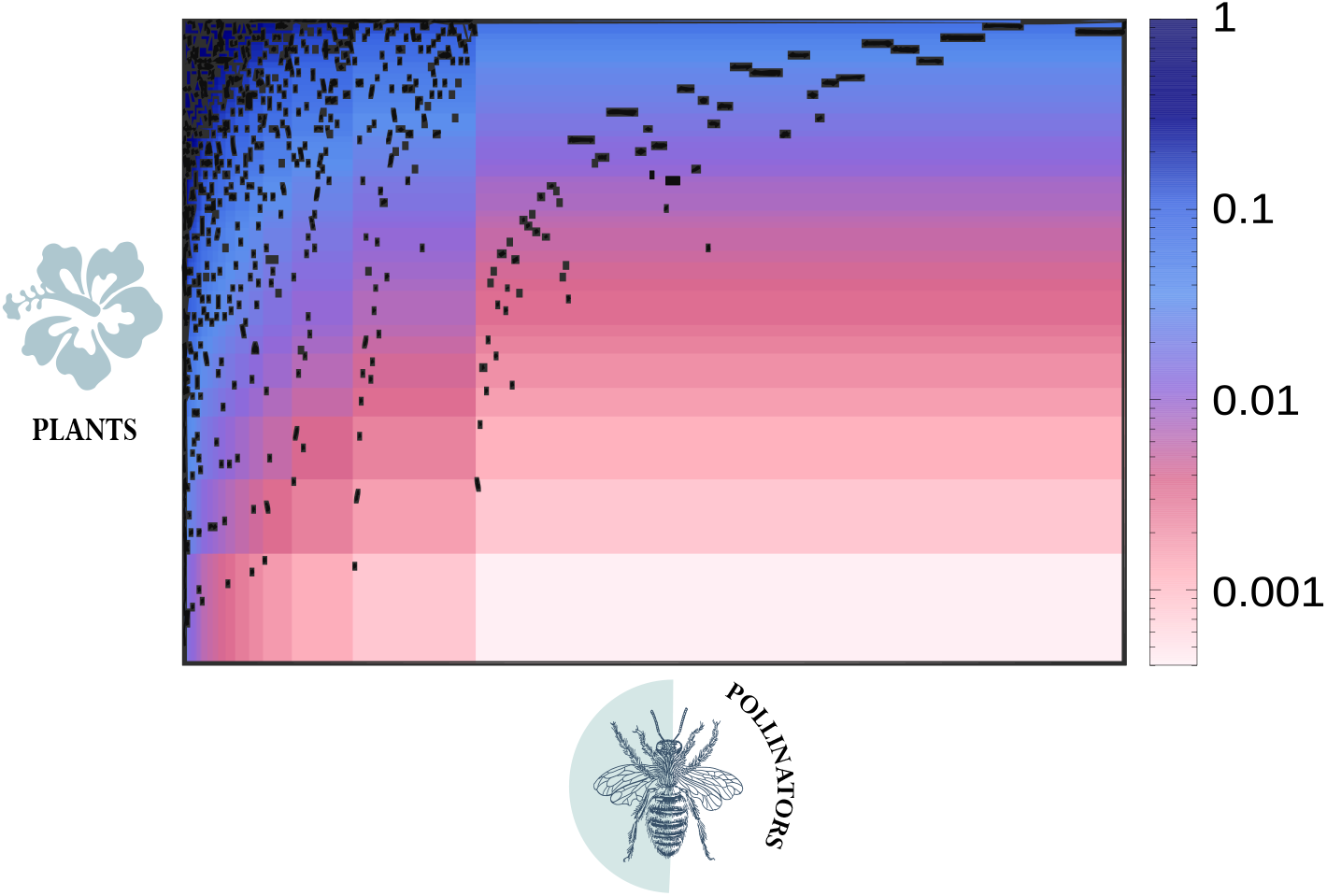
Comparison of generated and empirical mutualistic interactions. Probability of interaction between species in the statistical ensemble (color coded as indicated), for the plant-pollinator network recorded by Inoue et al. [23]. The empirical corresponding bipartite matrix of interactions is superimposed in black. Both plants and pollinators species have been ordered in decreasing order of their degrees (from top to bottom and from left to right). As it can be seen at a glance, the obtained probabilities are consistent with the observed interactions, with the dark regions delimiting an upper left triangle, as in an ideally nested structure. Note that the color legend is in logarithmic scale.

For each one of the 167 empirical networks we have obtained the interaction probability between the elements of the corresponding bipartite matrix. These networks include three different kinds of mutualistic communities: plant-pollinator, seed-disperser and plant-ant (see Section SI5 of the *SM*). A comparison between the obtained mean expectations of the nestedness of the randomized ensembles and the measured values of the nestedness of the real networks unveils a striking agreement, see Fig. 2. As reported in Table 1, the absolute difference between these two quantities is less than *one* standard deviation for 100 out of 167 networks (59.9%), raising to 158 out of 167 networks (94.6%), if we account for *two* standard deviations. The previous percentage increases further after performing a multiple testing correction (see *Methods*): we find that only 3 out of the 167 empirical observations of nestedness are significant (*p*-value < 0.05). The three of them, which are of a relatively small size (≤ 55 species), were found to be less nested than predicted by the statistical ensemble. Additionally, in order to ensure that our findings are not an artifact of using the NODF metric, we have performed the same analysis by using the largest eigenvalue radius, which has been recently proposed as an alternative way of measuring nestedness [12]. In this case, since it is not possible to obtain an analytical and derivable expression of the metric, we broadly sampled the statistical ensemble using the calculated probabilities and performed the statistical measures on the obtained samples (see *Methods*). This supplementary analysis produces results that are in agreement with those reported above for the NODF, with only 16 out of 167 networks unexpectedly nested (see Section SI3 of the *SM*).

**Figure 2.**
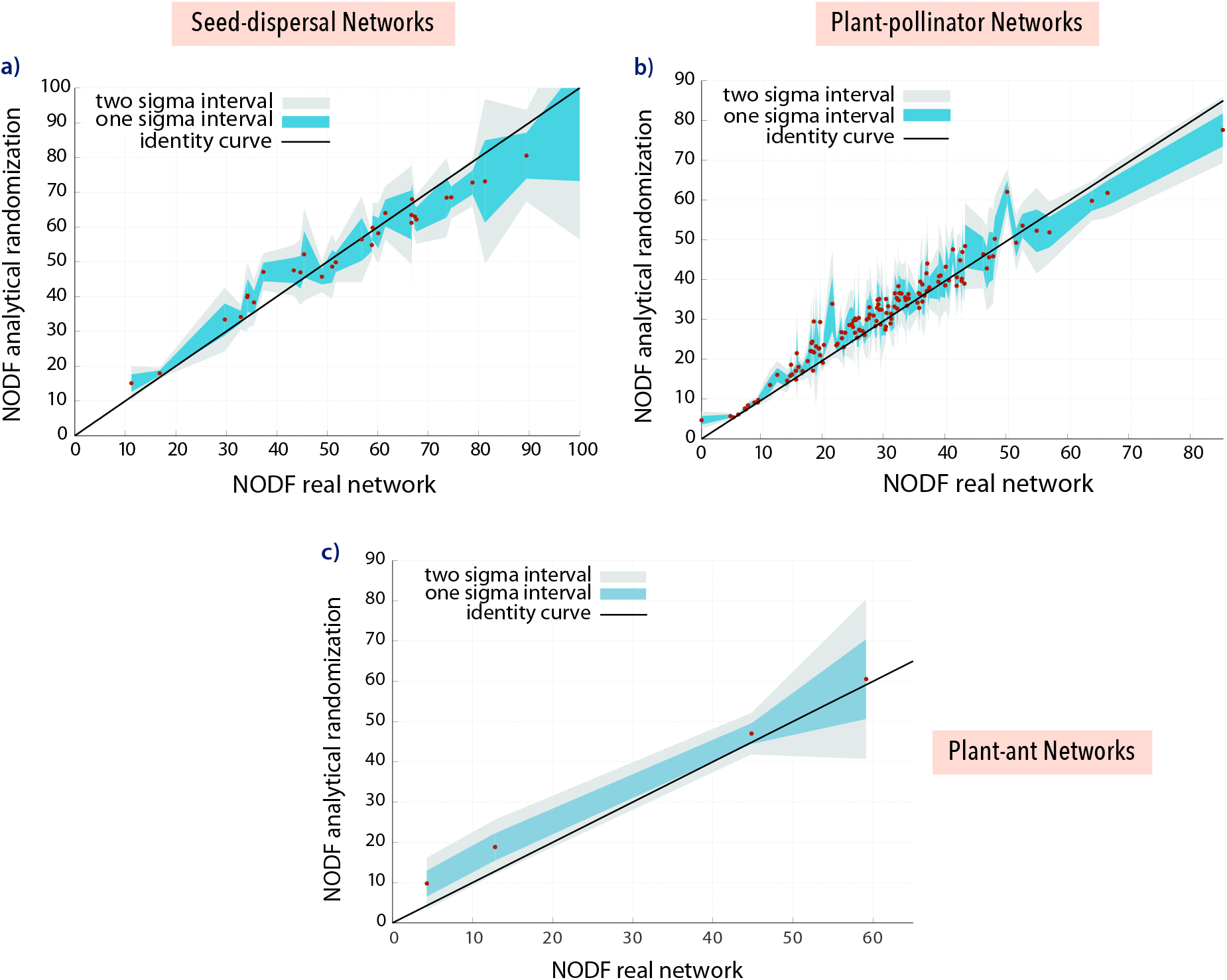
Determinants of nestedness. Panel (*a*): relative change in nestedness and the corresponding change in heterogeneity, measured for the set of 167 empirical networks and the average over the respective rewired ones. We used the rewiring algorithm described in *Methods*. Nestedness is measured using the NODF metric, whereas the heterogeneity is measured through the variance of the degree sequence of the unipartite adjacency matrix. We found a correlation index for a linear fit (excluding the top outlier) of R = 0.88. This closely linear relationship discovers a tight bound between nestedness and heterogeneity. Panel (*b*) shows a comparison between the real observation of the degree assortativity *r* (Pearson’s coefficient among degrees) and the average estimation in the statistical ensemble, for the 167 networks of our study. The fact that *r* < 0 for all values indicate that both real networks and the average of the randomized ensemble are naturally disassortative.

**Table 1.**
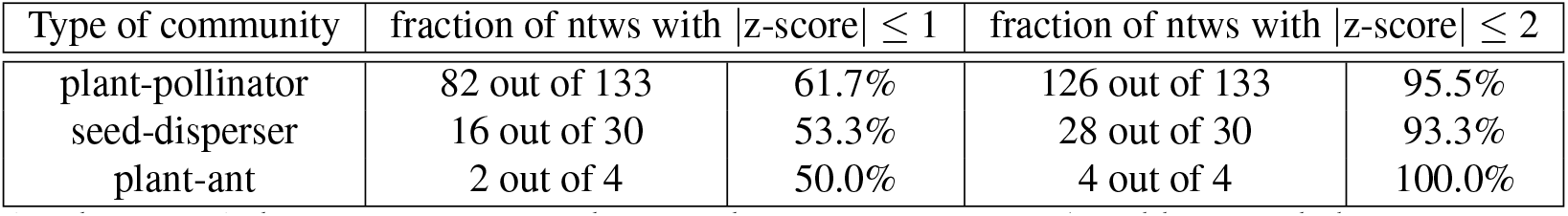
Results, disentangled into communities, showing the fraction of networks (abbreviated above as ‘ntws’) whose discrepancy between the real and randomized nestedness is less or equal than one or two standard deviations.

The findings above are of utmost importance in at least two fundamental aspects. Firstly, they demonstrate that, given the degree sequence of real networks, the observed nestedness is not significant. Secondly, they show that nestedness is not an irreducible pattern, in sharp contrast to the widely extended belief that it is so. In other words, these results reveal that the observed nested structure of the ecological communities studied is, in fact, a mere consequence of the degree sequences of the two guilds. Moreover, regarding recent debates about the use of a proper null model for nested networks [24], our findings point out the need of incorporating the information contained in the degree sequences. Indeed, our results indicate that an appropriate null model is the set of exponential random graphs for which the probability of having the same degree sequences of both branches of the bipartite graphs when compared to a real mutualistic network is maximized. Thus, we propose that the methodology implemented here to obtain the statistical ensemble of graphs that are compatible with the real networks could be a general tool to assess nestedness’ significance.

In the light of the previous results, the second question arising is whether we can determine which characteristic of the degree sequences modulates how nested a network is. Considering that the degree distributions of mutualistic communities have been reported to commonly follow a (truncated) power-law [25], we propose, as a plausible candidate, the *heterogeneity* in the number of contacts per species. Thus, our hypothesis is that for two networks with identical number of species and connections but diverse degree sequences, the most heterogeneous one (taking into account both guilds) will be as well the most nested. To evaluate this conjecture, we made use of a self-organizing network model that is devised with the aim of optimizing the nestedness of a network [5] by rewiring existing links (see *Methods*). After applying this algorithm to our empirical set of networks, we found that the resulting degree sequences are, with respect to the original ones, more heterogeneous and that the final networks are more nested, see Fig. 3a. This allows to bridge the gap between two topological features that have been classically treated separately, though previous works already suggested their connection [11,14]. Interestingly enough, the relationship between network’s heterogeneity and nestedness also explains why dynamical implications once attributed to nestedness like the sustainability of communities with a large number of different coexisting species [2] or the network’s structural stability [3,26], have recently been successfully associated with other properties such as the heterogeneity itself [15] or the species’ degree [13].

**Figure 3.**
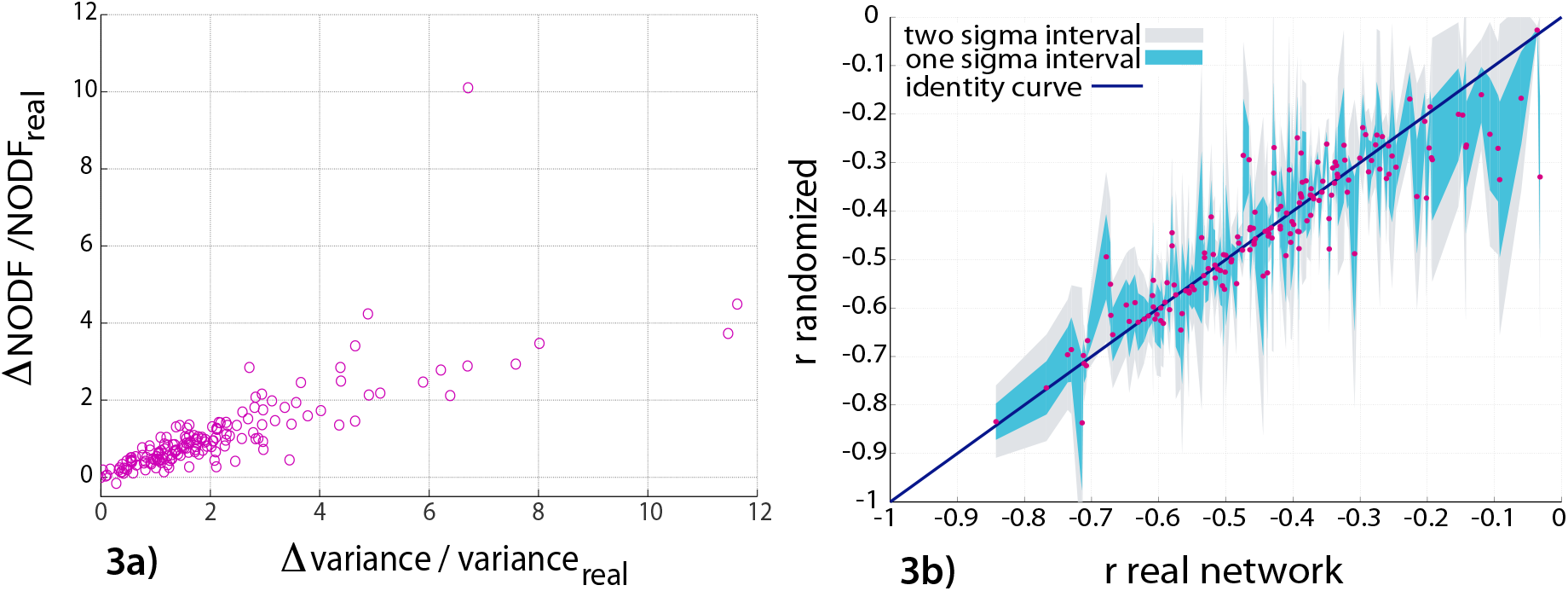
Determinants of nestedness. Panel (*a*): relative change in nestedness and the corresponding change in heterogeneity, measured for the set of 167 empirical networks and the average over the respective rewired ones. We used the rewiring algorithm described in *Methods*. Nestedness is measured using the NODF metric, whereas the heterogeneity is measured through the variance of the degree sequence of the unipartite adjacency matrix. We found a correlation index for a linear fit (excluding the top outlier) of *R* = 0.88. This closely linear relationship discovers a tight bound between nestedness and heterogeneity. Panel (*b*) shows a comparison between the real observation of the degree assortativity *r* (Pearson’s coefficient among degrees) and the average estimation in the statistical ensemble, for the 167 networks of our study. The fact that *r* < 0 for all values indicate that both real networks and the average of the randomized ensemble are naturally disassortative.

Moreover, accounting for the heterogeneity offers some further insight on the process of emergence of nestedness out of the degree sequences. At first glance, it might not be evident why our null model reproduces so well the empirical nestedness. A priori, we would naively expect that the random ensemble contains both nested and non-nested structures alike, in which specialists appear attached, respectively, to generalists or to other specialists. Although a given number of connections are certainly imposed by the existence of super-generalists as well as by finite size effects, normally there is still room for reshuffling links (like in the "swapping algorithm" [27]). In terms of mixing, we would say that, concerning specialists, both assortative configurations (nodes have neighbors with degrees similar to their own) and disassortative ones (neighbors have dissimilar degree) are in theory feasible. Why, then, our algorithm is expected to generate disassortative networks (see Fig. 3b)? Here, the particularity that we used a maximally-entropic ensemble plays a crucial role. Johnson et al. [28] showed that, in the case of heterogeneous systems, disassortativity is generally more entropic, that is, it is more likely as long as no external pressures are at work. This occurs, to put it simply, because for a species with few interactions there exist many more chances to engage with another species with numerous connections than matching to a low-connected partner. Therefore, the low significance of empirical nested patterns reported here is directly related to the fact that the number of mutualistic interactions per species is a highly heterogeneous quantity.

In concluding, it is worth mentioning that in recent years, nestedness has been proposed to arise either as an ecological feature that provides an optimal balance between competition and mutualism [2,6], or as a byproduct of processes such as the assembling rules [7,29]. Our results imply that no selective pressure has acted upon nestedness, which does not exclude, however, that such pressure has shaped the degree sequences. Even though such conclusions do not invalidate nestedness’ usefulness as an indicator of stability or robustness, we would like to underline that our findings clearly demonstrate that the degree sequences are the lower-order determinants of nestedness. Moreover, this highlights the interest of focusing on the ecological and evolutionary mechanisms that have led to the heterogeneous degree distributions present in mutualistic ecosystems [30, 31], like might be the need to diminish the cost of mutualism [32]. Understanding the way in which structural properties emerge in ecological communities is a fundamental, long-standing challenge that can provide critical clues to depict ecosystems’ past assembling, present functioning and future responses. Finally, given that nested patterns have been recurrently detected across systems as diverse as biological, social and technological networks, our findings are expected to have relevant implications beyond the present analysis of ecological mutualistic communities.

## Methods

### Construction of the Random Ensemble

We constructed an ensemble following the Exponential Random Graph model. This ensemble maximizes the Shannon-Gibbs Entropy given the average degree sequences of the two guilds of a bipartite network as constraint. Yet, it is not fully determined due to the presence of some free Lagrange multipliers resulting from the constrained optimization. Following Squartini and Garlaschelli [20] [22], we imposed that the degree sequences of the empirical network are found with maximum likelihood. This provides a set of coupled equations to solve for the Lagrange multipliers (one equation per node, see Eqs. 7-8 in Section SI1 of the *SM*). Determining the statistical random ensembles of our 167 empirical networks entails solving computationally 167 optimization problems. For each network, we numerically found the Lagrange multipliers that maximize the likelihood using two different, independent algorithms: *I*) a global, pseudo-random numerical method for optimizing the likelihood and *2)* a deterministic, gradient-based algorithm for solving non-linear systems of equations. See Section SI1 of the *SM* for the explicit expressions of the ensemble probability and equations to solve, as well as more information on the numerical implementation. As shown in [20], in the case of local constraints (as the degree sequences), the probability of existence of a graph in the ensemble can be exactly factorized into the probabilities of existence of a link between species [20]. Therefore, after numerically determining each optimal set of Lagrange multipliers, we built the matrix containing the average probability of interaction corresponding to each empirical network (see an example in Fig. 1 and Eqs. 9-10 in Section SI1 of the *SM*).

### Statistical Measures on the Random Ensemble

We performed the statistical measures on the ensemble following either of the two following approaches. On the one hand, as long as the property that we aim to evaluate could be formulated as an analytical and derivable expression, Squartini and Garlaschelli showed [20] that it is possible to obtain, at first order, the analytical expression of the first and second moments of the corresponding distribution. These expressions depend only on the link probabilities (see Eq. 1–2 in Section SI2 of the *SM*). We wrote the NODF metric in a compact, analytical form and derived the expression of the theoretical average expectation and standard deviation of nestedness in the ensemble (Eqs. 7-9 in Section SI2 of the *SM*). Finally, we obtained the 167 probability matrices of interactions and computed the main statistical moments. On the other hand, one can always sample the ensemble in order to study the statistics of the target property on a generated sampling. Using this scheme, we produced 10^4^ networks that were assembled using the obtained probability matrix of interactions. Over this subset, we numerically calculated the average expectation and the standard deviation of the largest eigenvalue radius [12] (see Section SI3 of the *SM*) and the assortativity index measured through the Pearson coefficient of the degrees (see Section SI4 of the *SM*).

### Significance tests

We quantified the significance of the nestedness using the *z-score* index, which for a general property *x* reads: 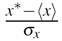. For us, 〈*x*〉is the average nestedness computed in the ensemble, and we compare it with the empirical observations *x*.* The standard deviation is *σ*_*x*_. Given that the NODF values are gaussian distributed in the random ensemble (see Section SI2 of the *SM*), the *z*-scores can be directly related to *p*-values. We performed a *multiple test correction* which allows accounting for the fact that as the number of statistical tests increases, so does the probability of finding rare events [30]. Thus, when considering the multiple comparisons we could prevent overstating the number of significant discoveries. It is pertinent to apply this technique here since the 167 cases studied are evaluated under the same *null hypothesis* and all of them follow a normal distribution. We employed the *false discovery rate* method, in particular the Benjamini-Hochberg procedure which applies to independent tests [30].

### Self organizing network model

In order to reorganize the original network into an even more nested structure, we numerically implemented the self-organizing network model proposed by Burgos et al. [5]. This methodology keeps constant many aspects susceptible to affect the measure of nestedness, like the size and fill, but modifies the degree sequences through the redistribution of connections. We rewired the links among species following two simple rules: *i)* when changing an interaction, the new partner must have higher degree than the old neighbor *ii)* if the proposed redistribution leaves one of the two nodes with no interactions at all, we reject the change. This operation was repeated until the system achieved a frozen state in which no more reconnections were accepted (we considered this happened when 10^3^*N* consecutive rejections occurred, being *N* the number of nodes of the network). The final frozen state is normally not perfectly nested, since condition *ii* typically leads to configurations which are not utterly optimal. To compensate this, we carried out 10^3^ independent rewiring operations for each network. We then averaged the target properties, namely, nestedness (measured using NODF) and the variance of the joint degree sequence of the two guilds.

## SUPPLEMENTARY INFORMATION

## S1. Construction of the random ensemble

This Section provides additional details on how we constructed the statistical ensembles. An ensemble is a set of networks across which unconstrained features will vary randomly, and over which we will perform statistical measures.

### General randomizing scheme

We denote a network in the ensemble by its graph **G**, except for the real network which is **G**\*. We characterize the ensemble by the probability of occurrence of each of its elements, ***P***(***G***). Following [31] and [20], we choose a probability such that the constraints are only satisfied on average, thus allowing slight mismatches across the ensemble. This is equivalent to constructing a *grand-canonical ensemble* (as opposite to the *micro-canonical ensemble,* where the constraints need to be exactly met always).

As proposed in [31], we ask the probability of each graph in the ensemble to maximize the Shannon-Gibbs entropy, defined as:

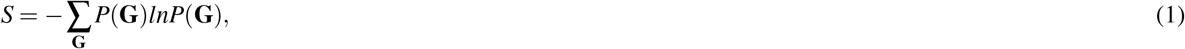

where the sum runs over all the graphs *G* in the ensemble. This leads to the *Exponential Random Graph model,* which reads:

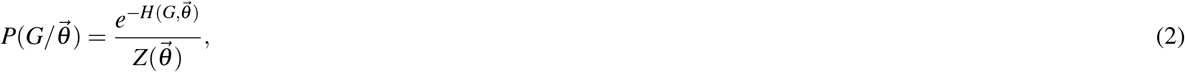

being *H* the graph Hamiltonian such that 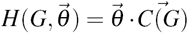, and Z the normalizing partition function 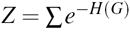. The set of variables 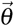 are the *Lagrange multipliers,* resulting from the maximization of Eq. 1 under the chosen constraints 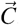.

Secondly, we proceed to calculate the exact values of the Lagrange multipliers. Following Squartini and Garlaschelli [20] [22], we determine these parameters by imposing that the real network is found in the ensemble with maximum probability. Indeed, we may write the log-likelihood of observing the real network 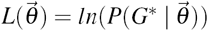 as:

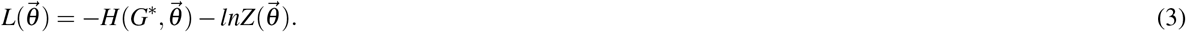

Maximizing this quantity thus allows fixing the 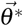 values. This second requirement ensures not only that the constraints are met on average, but also that they are the most likely ones, which is a warranty of non-bias [20].

### Ensemble for a bipartite network with constrained degree sequences

We now explain how the randomizing scheme by Squartini and Garlaschelli [20] applies to our specific problem, namely a *bipartite network* subject to *local* constraints. The scheme has already been applied to study international trade networks [21].

To begin with, we construct the hamiltonian for a bipartite network, whose bipartite matrix we call **B**. At variance with the monopartite case, we have *two* degree sequences (one for each of the guilds) which need to be taken into account separately. Although the scheme is equally valid for any mutualistic network (seed-dispersers, ant-plants…), for the sake of clarity we restrict our notation to the paradigmatic case of plant-pollinator communities. Thus, we will speak of systems with *N_p_* number *of plants* and *N*_*A*_ pollinating *animals.* The constrained degree sequences, given by the real network, will be represented respectively by 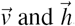, where *v*_*p*_ is the diversity of *visiting* animal species that a plant species *p* receives, while *h*_*a*_ is the number of different *hosting* plant species with which a pollinator species *a* interacts.

In order to enforce both distributions as constraints we introduce two sets of Lagrange multipliers, 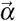 for plants and 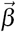 for animals. Subsequently, the graph hamiltonian can be written as

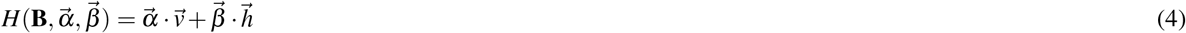

This means that the probability, Eq. 2, of encountering a bipartite graph *B* in the exponential random graph ensemble becomes:

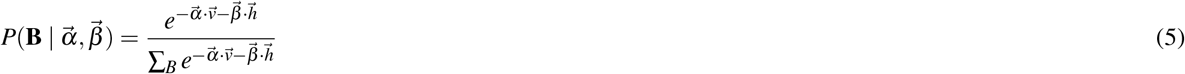

To simplify the notation we introduce the variable change 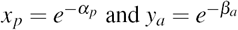, as suggested as well by Squartini and Garlaschelli. Then, the log-likelihood of encountering the real network is:

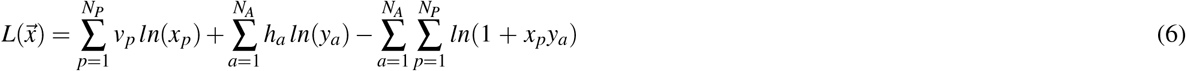

which we need to maximize in order to find the optimal variables 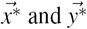, that ultimately define our ensemble. Indeed, by requiring that 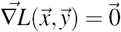, we obtain the following set of equations:

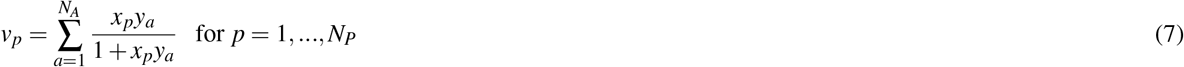

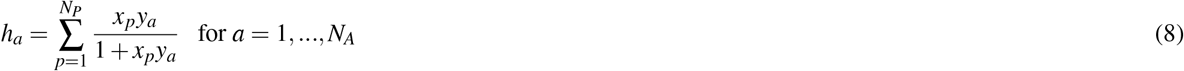

It can be easily shown that these equations are equivalent to imposing that the average degrees (right hand side) are equal to the degree sequence from the real network (left hand side), as we do below.

### Probability matrix of interactions

Garlaschelli and Squartini also showed [20] that in the case of local constraints, the ensemble probability can be factorized, using our notation, in terms of the *probability of existence of a link between a plant species ‘p’ and animal species ‘a*’, which we call *p*_*pa*_. In effect, by taking 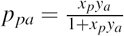, replacing it into equation 5 and doing some little algebra, one finds:

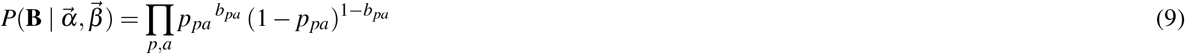

Where *b*_*pa*_ is the (*p,a*) element of the bipartite matrix of interactions. Then, using expression 9, it is almost immediate to see that 〈*b*_*pa*_〉= *p*_*pa*_, thus in turn, 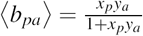. This shows that, as we had said, the right hand-side of equations 7-8 is a sum over a column or row of expected values of the randomized bipartite matrix.

We also note that the possibility of factorizing 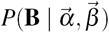 essentially entails that the probabilities *p*_*pa*_ are independent among them. In other words, when the constraints enforced are local, the probability of existence of different links are independent among them. This automatically allows the construction of the exact expected randomized matrix of interactions:

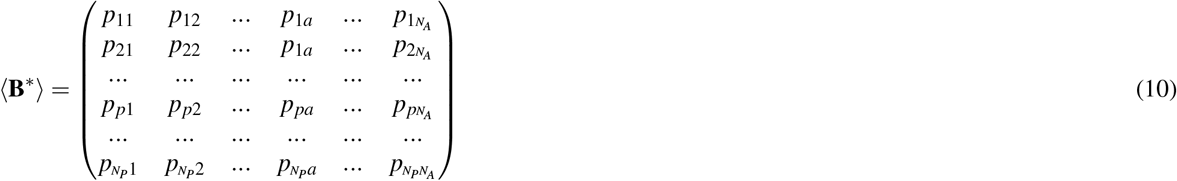

### Computational implementation

Here we give the numerical details on how we obtained the Lagrange multipliers 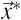 and 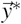 that eventually define the corresponding statistical ensembles of the empirical networks. As proposed by Squartini and Garlaschelli [20], encountering these multipliers might be achieved following either of two procedures: by directly maximizing the log-likelihood in Eq. 6 through an optimizing algorithm, or by solving the non-linear, coupled set of equations in 7-8.

First, we optimized the log-likelihood by means of a global search, pseudo-random algorithm belonging to the Monte-Carlo family and known as *simulated annealing* [35-37]. In short, this method aims to find the global minimum of a function by sequentially exploring the solution space, through producing random proposals subjected to an accepting criteria. More specifically, it uses the Metropolis criteria [38], which is driven by a parameter *T* traditionally called *temperature.* The algorithm works iteratively, by finding the most probable state at each temperature and then using it as the initial condition in the next step. We start with a high temperature, which means that almost all proposals are accepted. As the algorithm progresses, the temperature decreases and so does the acceptance probability. This makes the search to become more and more restricted around eventual solutions, until a certain number of consecutive iterations (in our case, five) have produced solutions differing less than a certain tolerance, called *tol.* When that happens, we are confident enough of having reached the ground state and the algorithm stops. Given the pseudo-aleatory character of this approach, which allows escaping from local hills, it is extendedly used in situations in which the co-existence of several local optima is suspected.

In our case, since the algorithm is originally intended for minimizing but we aim to maximize, we simply took the minus of the function. Additionally, instead of optimizing the log-likelihood as written in Eq. 6, we incorporated the fact that the degrees may be degenerate. This means that nodes of the same guild having identical degrees satisfy equivalent equations, hence necessarily bearing the same solution. To account for this, we introduced a multiplicity factor *m*_*p*_ for plants and *m*_*a*_ for animals. If we call *red*_*P*_ and *red*_*A*_ the redundancy for plants and for animals (namely, the corresponding numbers of repeated degrees), then the system can be redimensionalized to 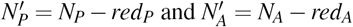. Consequently the log-likelihood might be rewritten into:

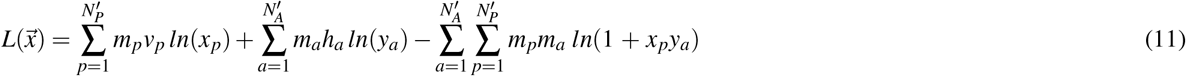

Although, in analytical terms, the original expression in Eq. 6 and this latter one are obviously equivalent, from a computational point of view reducing the number of variables enhances the algorithm’s efficiency. Besides, imposing from the beginning such identity between variables improves the accuracy of the program.

We programmed a classical version of simulated annealing, with a starting temperature of *T =* 10^3^, a reduction factor of the temperature of *RT =* 0.85, a tolerance *tol =* 10^-6^ and a total number of updates per fixed temperature of 2·10^4^. Furthermore, we ran the algorithm 10 times per network with different random seeds, in order to produce independent sequences of explorations. When all runs convergence to the same solution, as outlined by Goffe [37], it is extremely probable that we have certainly encountered the global optimum.

Secondly, we solved the set of equations by means of a local, deterministic algorithm known as the *modified Powell hybrid method.* In particular, we used the MINPACK library [39] for FORTRAN, available online [40]. This method finds the zero of a non-linear system by exploiting its Jacobian, which we analytically calculated and implemented into the program.

Like before, we re-dimensionalized the problem to 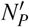 equations for plants and 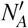 equations for animals, which now read:

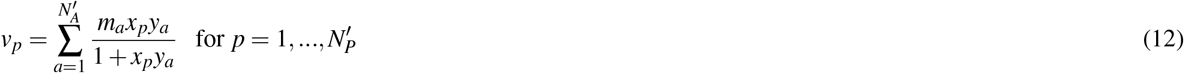

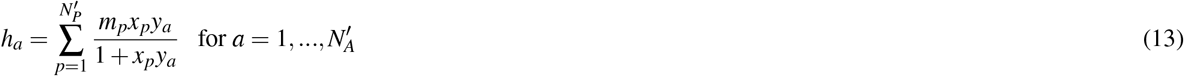

We implemented these equations and their Jacobian and ran the algorithm with a tolerance *tol =* 10^-11^ (as defined in the source code). The possibility of exploiting the gradient provides, in general, a greater local accuracy than the simulated annealing technique. However, its shortcoming lays in the risk of getting trapped in local optima, from which, due to its deterministic nature, it is unable to escape. To compensate this drawback we performed a significant sampling of the space of initial conditions, by running 10^4^ iterations of the algorithm, each with a different random selection of starting points, covering as well distinct ranges. However, due to the encounter of rough, rather accidental configuration surfaces, the modified Powell hybrid method was not always able to converge to a solution. The rate of success was approximately 50%.

To finally ensure that we found the global maximum, we compared the outcomes of the various independent runs and also, when the Powell algorithm succeeded, among both methods (so in total 10 runs for the simulated annealing and 10^4^ for the Powell hybrid method). We confirmed, in all cases, that the same maxima was found. This enables us to assume that we found the actual *global* optimizing Lagrange Multipliers for each one of the networks of our study.

Moreover, the constraints were successfully met with a relative precision between 0.01% and 10%. This check was carried on by computing the average degrees using equations 7-8 and comparing the output with the imposed degree sequences (extracted from the empirical networks). The worst case of 10% was typically caused by discrepancies in low degrees, generally the most sensitive to imprecisions in the elements of the randomized matrix (since the matrix elements of low degree nodes are usually very small, see Fig.1 in main text as an example). Altogether, this second check warrants that our constrained optimization worked as aimed.

## S2. Nestedness statistical measures using *NODF*

Here we describe how to obtain statistical measures in the random ensemble through analytical expressions, and particularly present our derivation for the nestedness metric known as NODF [41].

### General analytical expressions

Let us call a property by *X* and its randomized measure (that is, the average across the random ensemble) by 〈*X*〉*. When the property *X* can be calculated through an *analytical* expression (non-algorithmic) as a function of the bipartite matrix **B**, then Squartini and Garlaschelli [20] showed that it is possible to perform an approximate but accurate measure of the first and second moments of *X, directly* on 〈**B\***〉 (see Eq. 10, SI1). In particular, for the bipartite case, this reads:

**Figure 4.**
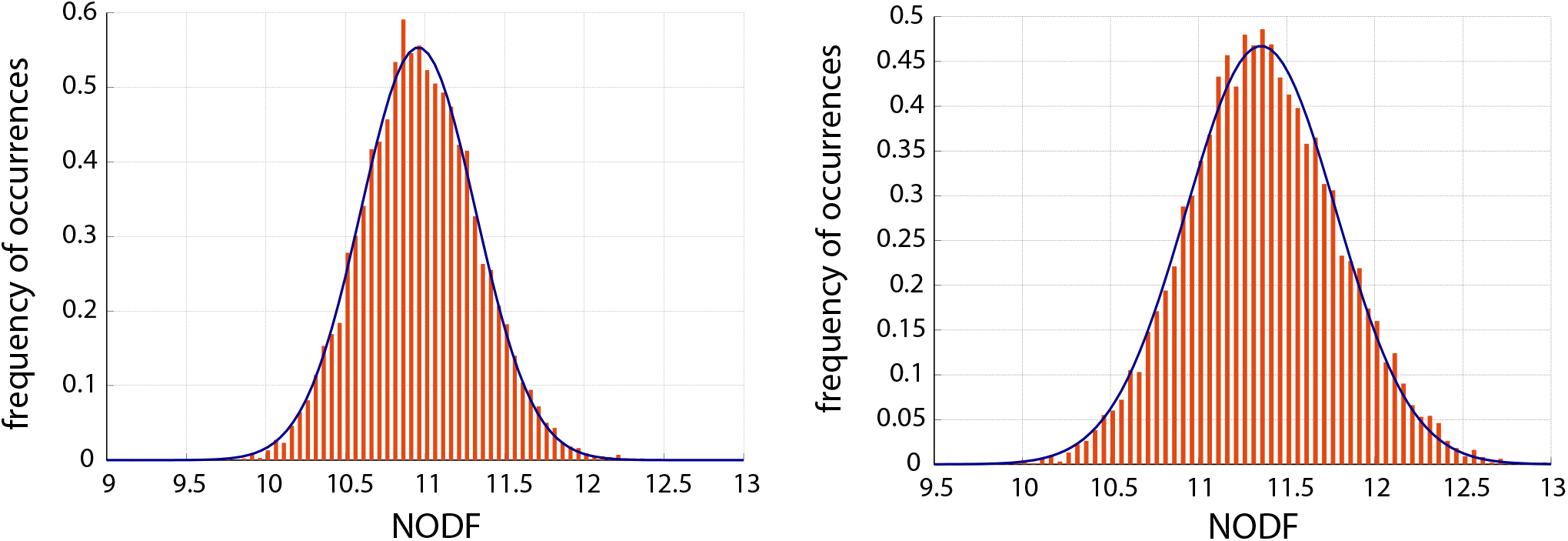
Nestedness distribution for two samplings of the statistical ensembles corresponding to the empirical networks by Petanidou et al. [42] (left) and by Inoue et al. [23] (right). In blue, fits of a gaussian function using the mean and standard deviation extracted from each distribution.

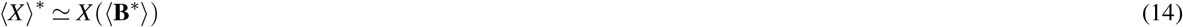

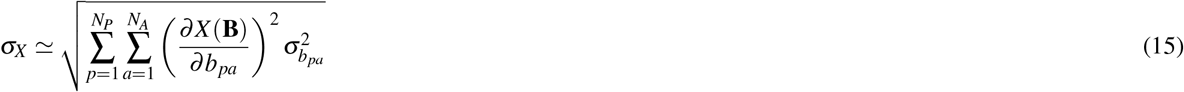

Where *σb*_*pa*_ is the standard deviation for the bipartite matrix element *b*_*pa*_. The condition for these approximations to be accurate is that the property *X* be gaussian-distributed in the random ensemble.

### Derivation for *NODF*

Let us see how the previous expressions can be applied to measure nestedness with the *nestedness metric based on overlap and decreasing fill* (hereafter, *NODF)* by Almeida-Neto et al. [38].

We first verified that the assumption of gaussianity is fulfilled by performing a check on a smaller subset of our set of empirical networks. To do this, for each of the corresponding statistical ensembles we generated a sample of 10^4^ networks obeying the probability of link existence given by 〈**B**\*〉 (see Eq. 10, SI1). We then computed the nestedness of each sampled network in order to generate the nestedness distribution. In all cases we could successfully fit a gaussian function (see Fig. 5 as an example).

**Figure 5.**
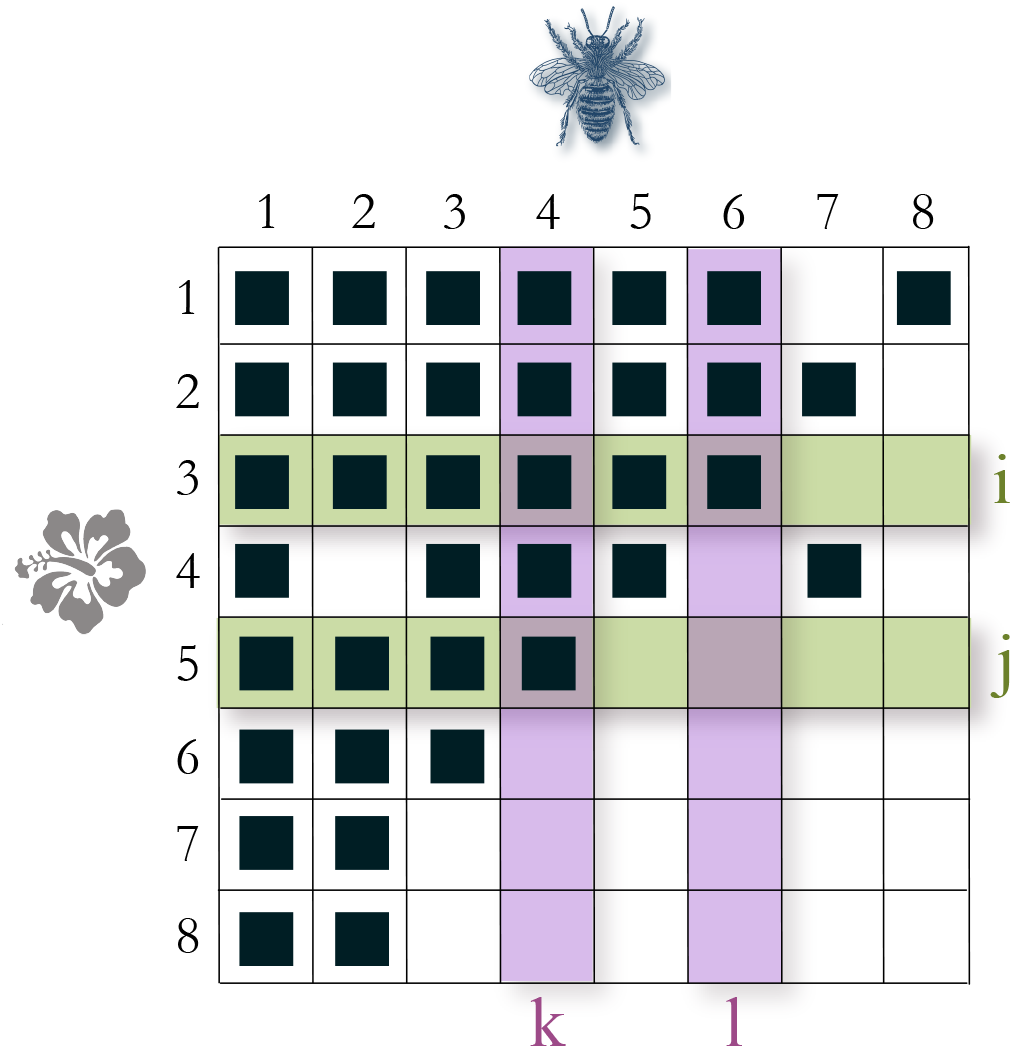
Example of an ordered matrix of interactions, not perfectly nested. Species of both guilds have been ordered in decreasing degree, and the numbered labels indicate their rank (the larger the degree, the smaller the rank). The indexes *i*, *j*, *k* and *l* illustrate our notation for rows and columns.

Once shown that the gaussian behavior is satisfied for this metric, we next apply Eqs. 14-15. Yet, we need an analytical, packed expression that facilitates the calculations of the metric. The NODF basically considers two contributing factors to nestedness: *decreasing fill* (the fact that, after a proper ordering, both degree sequences progressively decline) and *paired overlap* (the number of shared partners between two columns or rows, normalized by the smaller degree). By gathering together the sequential analysis indicated by Almeida-Neto et al., we proposed a novel compact expression to calculate NODF, that reads:

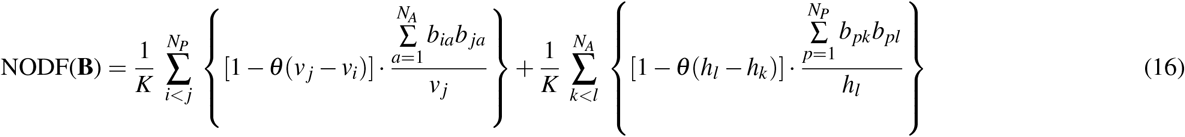

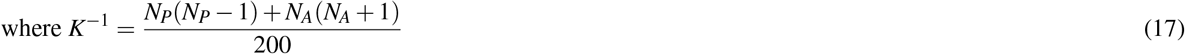

We maintain here our previous notation (see SI1), so *v*_*p*_ is the degree of plant *p* and *h*_*a*_ the degree of animal *a*. The double sums run over two indices such that, as seen in Fig. 5, row *i* is placed upper row *j* and column *k* more to the left than column *l*. The *K* factor contains the normalization over all possible pairs, as well as a percentage rescaling (note that NODF takes values between 0 and 100). Finally, the *θ* term is the Heaviside step function, which is zero when its argument is negative, and one if its argument is positive or zero. In our context it serves to encapsulate the decreasing fill condition. In fact, from now on we will use the following abbreviations:

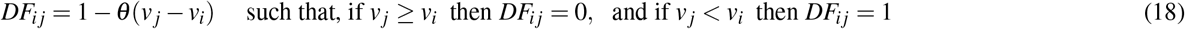

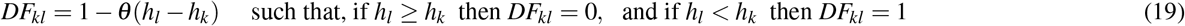

The analytical and packed expression for NODF that appears in Eq. 16 can then be plugged into Eqs. 14-15. Accordingly, we obtained that the first moment of the randomized NODF for a certain real bipartite matrix **B**\* reads:

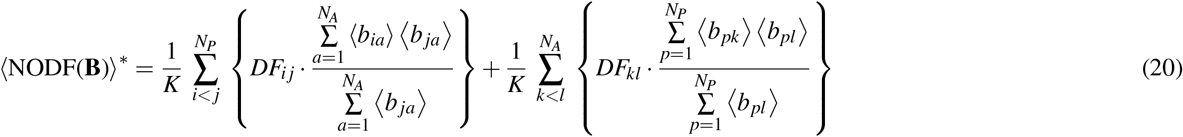

Notice that 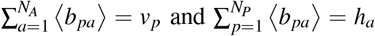, given that the randomized matrix necessarily fulfills the enforced constrains. Additionally, this warrants that the ordering of the matrix is equal to the original one, which is important since NODF is ordering-dependent through the decreasing fill terms.

It is also interesting to remark that the previous expression can be understood in probabilistic terms. Indeed, given that 〈*b*_*pa*_〉 = *p*_*ap*_, where *p*_*pa*_ are independent link probabilities, the overlap term might be seen as a joint probability of two independent events, divided by a normalizing factor which is the union of independent probabilities. For example, for one pair of animals, the overlap term results in:

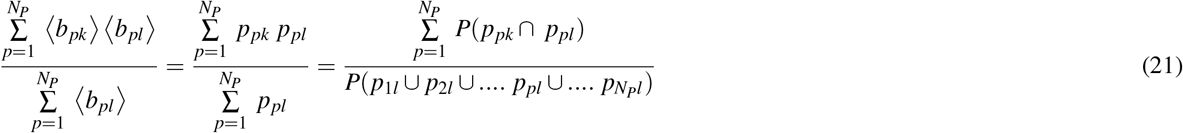

Apart from the first moment, equation 15 indicates how to compute the standard deviation. Adapting it to the nestedness measure, we encounter:

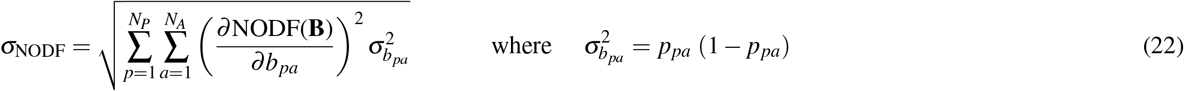

We have introduced that the standard deviation of the probability of links is that of a Bernoulli distribution, since the existence of a link is a binomial process (either there is interaction or there is not). Furthermore, the derivative with respect to a general matrix element *b*_*rc*_ (the sub-index *r* stands for rows and *c* stands for columns) can be split into the contributions of plants and of animals:

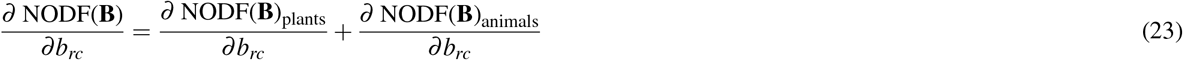

After deriving, we obtained that:

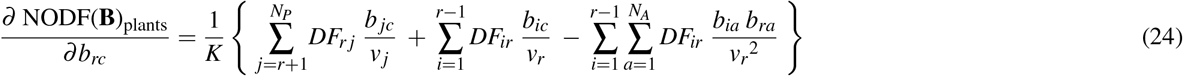

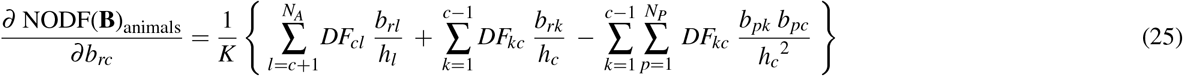

These novel estimations of the first and second moments can then be used to provide a randomized measure of nestedness 〈NODF(**B**)〉* together with its statistical significance.

## S3. Nestedness statistical measures using the *largest eigenvalue radius*

The largest eigenvalue radius was recently proposed by Staniczenko et al. [12] as an alternative measure for nestedness that directly relies on the spectral properties of the adjacency matrix. The fact that it involves finding the *maximum* eigenvalue entails that we lack an analytical and derivable expression for it. This, in turn, means that we can not derive for this metric the analytical expressions of Eqs. 14-15 in SI2. Therefore, in order to calculate the statistical properties of our interest, we produced a sample of the statistical ensemble and algorithmically computed the distribution of the largest eigenvalue radius.

In detail, we produced 10^4^ networks, sampled using the link probabilities in 〈**B**\*〉 (see Eq. 10, SI1). Then we computed the largest eigenvalue radius, which we call *ρ*(*λ*), using the *R* package *rARPACK* [43]. Finally we calculated the average and the standard deviation of the resulting distribution. As shown in Fig. 6, the results are fully compatible with those found using the NODF nestedness metric. See also Tables. 2 and 3 for details of percentages.

**Figure 6.**
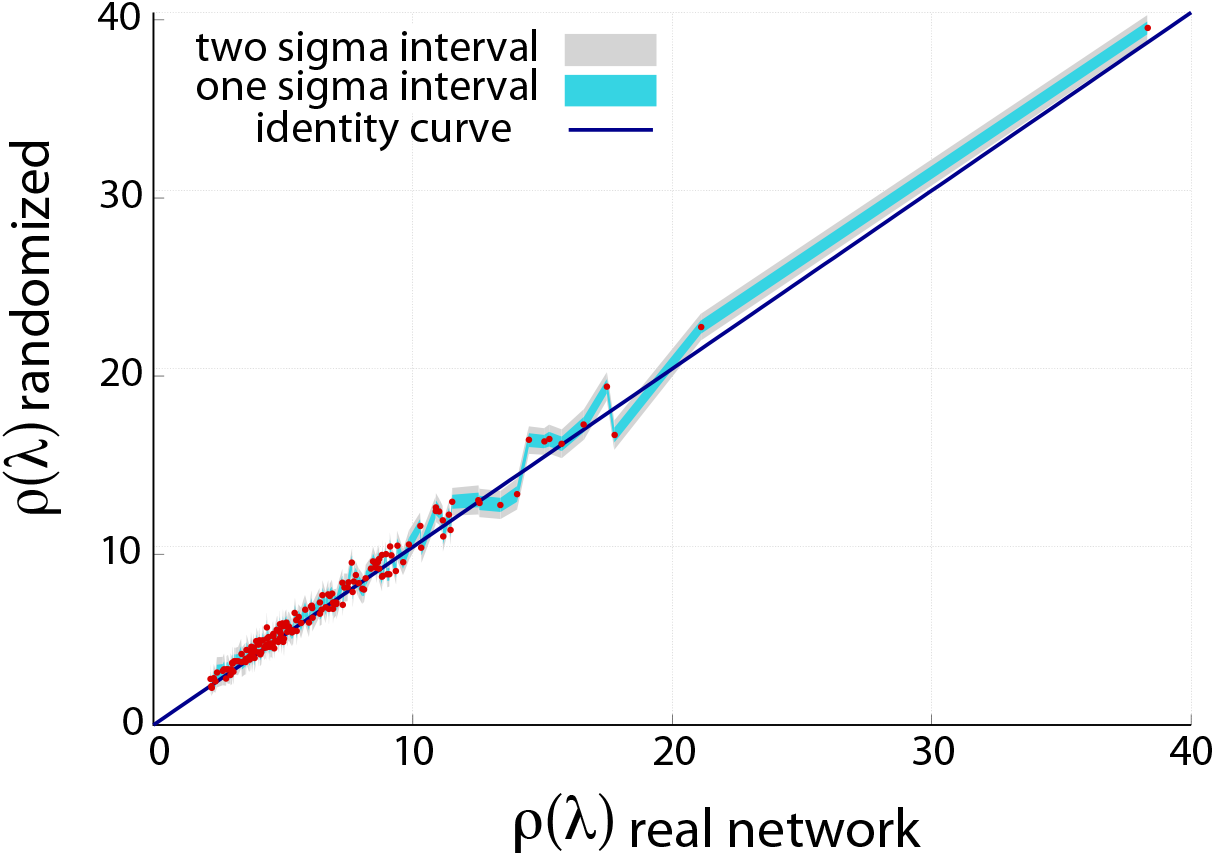
Comparison between the real observation of the largest eigenvalue radius *ρ*(*λ*) and the average estimation in the statistical ensemble, for the 167 networks of our study.

**Table 2.**
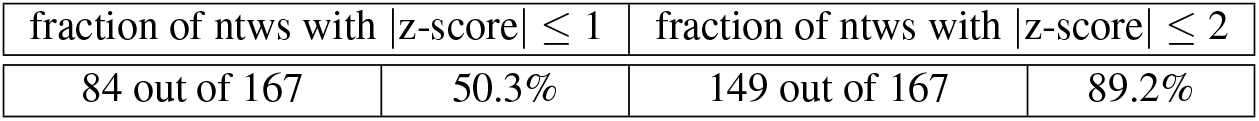
Fraction of networks whose discrepancy between the real and randomized nestedness is less or equal than one or two sigma.

**Table 3.**
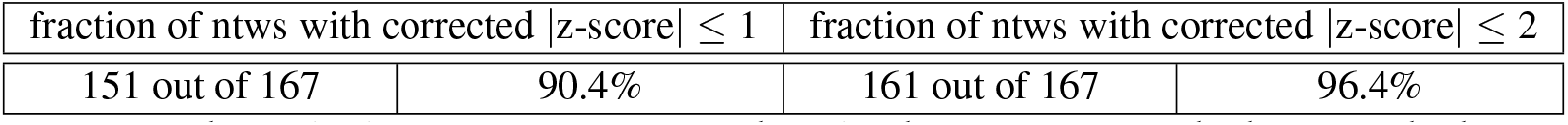
After performing the multiple test correction using the false discovery rate method (see Methods summary in Main text), fraction of networks whose discrepancy between the real and randomized nestedness is less or equal than one or two sigma.

**Figure 7.**
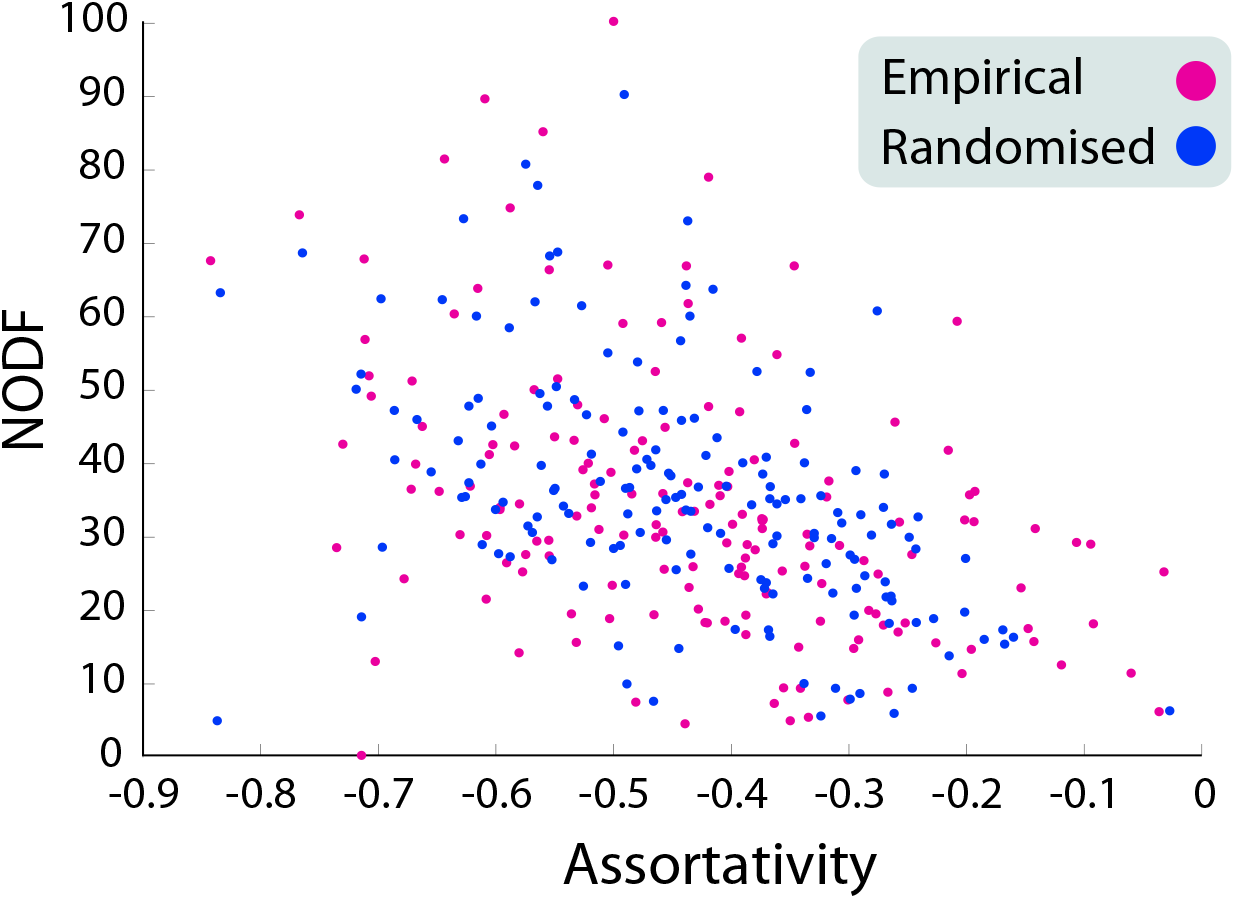
Relation between the degree assortativity and the nestedness for the real network (in red) and the average measure for the randomised case (in blue).

## S4. Assortativity statistical measures

Assortativity is a network feature that quantifies to what extent nodes tend to match other nodes that are similar (or dissimilar) to them. Here, we particularly used the notion of degree assortativity, which means that *similarity* is labelled by the degree. We followed the definition proposed by Newman [44], which consists of a normalized correlation coefficient between degrees. This eventually corresponds to the *Pearson correlation coefficient* denoted by *r*, such that *r* = −1 indicates perfect disassortativity, *r* = 0 no correlation at all and *r* = 1 maximum assortativity.

In order to compute the statistical properties of this quantity, we produced for each ensemble a sampling made up by 10^4^ networks. We then measured computationally the assortativity of each sampled network using the *assortativity_degree* function from the *igraph* package in *R* [45]. Finally this allowed us to calculate the first and second moments of the assortativity for each ensemble in our set.

## S5. Data

In our study we analyzed 167 real interaction networks from the *Web of Life* dataset [46]. This set consists of 133 plant-pollinator communities [references from 47 to 83], 30 seed-dispersal [references from 84 to 104] and 4 plant-ant [references from 105 to 108].

Data sometimes included information about link’s weight, but we converted all networks to binary matrices.

